## Supplementary Materials for "Re-focusing visual working memory during expected and unexpected memory tests"

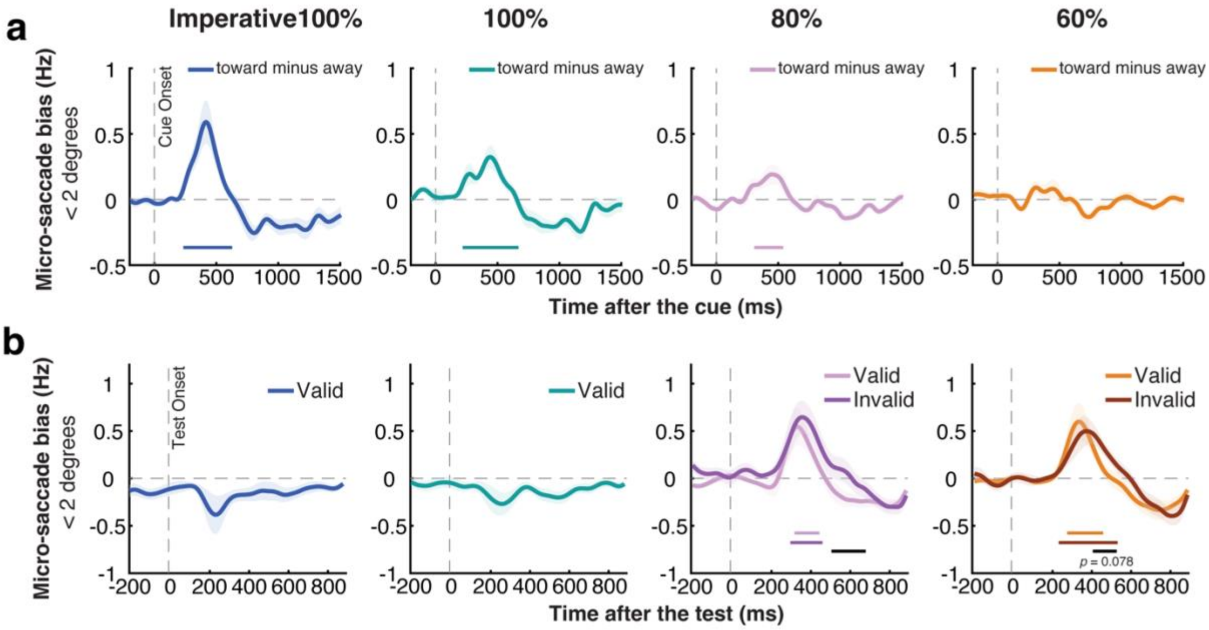


**Figure S1. Spatial saccade biases associated with initial orienting after the retrocue and re-orienting after the memory test are preserved when exclusively considering fixational "micro" saccades < 2 degrees. a**) Spatial saccade bias (toward minus away, calculated by exclusively considering saccades within 2 visual degrees) across conditions following the cue. **b**) Same plots as in **a** but following the memory test. Colored horizontal lines above x-axis indicate significant clusters. The black horizontal lines above the x-axis in panel **b** indicate significant difference clusters between validly cued (expected) and invalidly cued (unexpected) memory tests. Shading and error bars represent ±1SEM.

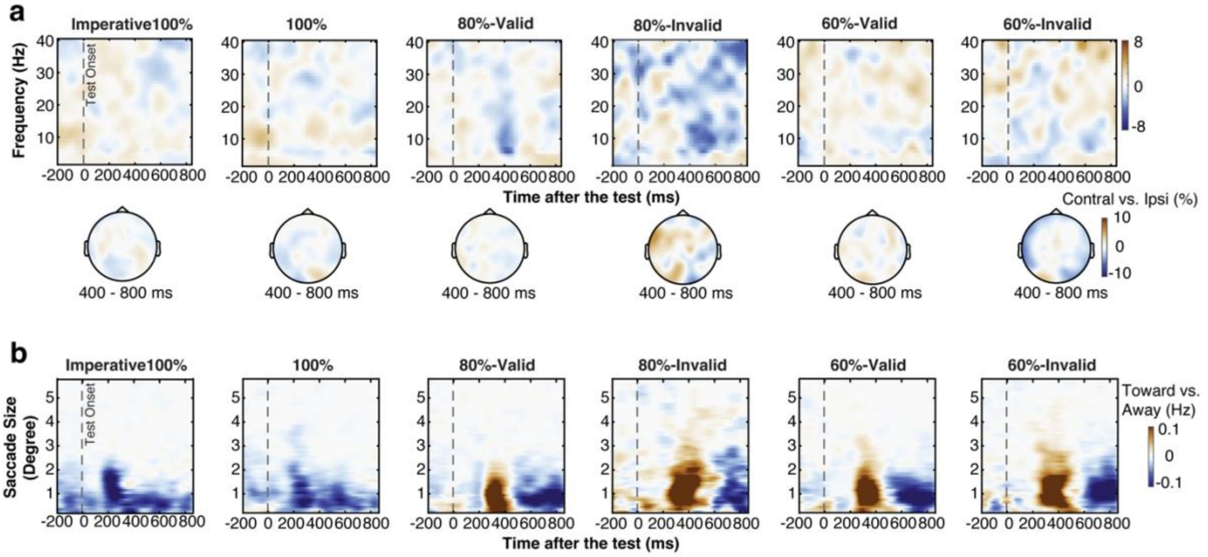


**Figure S2. Time-frequency and saccadic activities after the memory test. a**) Top: time-frequency spectrum (contralateral minus ipsilateral) across frequency bands (2-40 Hz) across conditions following the memory test. Bottom: topographic distribution of 8-12 Hz EEG-alpha lateralization across conditions (averaged across 400-800 ms after the test). **b**) Spatial saccade bias (toward minus away, color-coded) as a function of saccade size. The difference between toward and away saccades (with red colors denoting more toward saccades) was predominantly driven by saccades in the micro-saccades range (< 2°) rather than looking back to the original location of the items that were centred at 6 degrees during encoding.

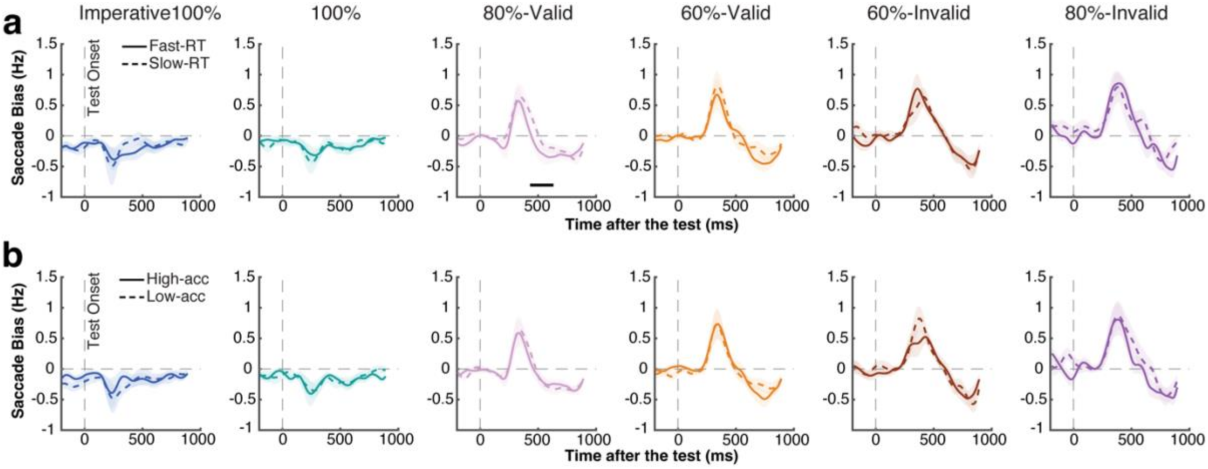


**Figure S3. Saccade biases associated with attentional re-orienting after the memory test as a function of behavioral performance. a**) Memory test-locked saccade bias comparison between fast- and slow-RT trials (median split) across cue-reliability conditions (color-coded). Solid and dashed time courses represent fast- and slow- RT trials, respectively. **b**) Similar comparison as in **a** between high- and low- accuracy trials (median split). Solid and dashed time courses represent high- and low- accuracy trials, respectively. The black horizontal line above the x-axis represents a significant difference cluster after permuation testing. Shading and error bars indicate ±1SEM.

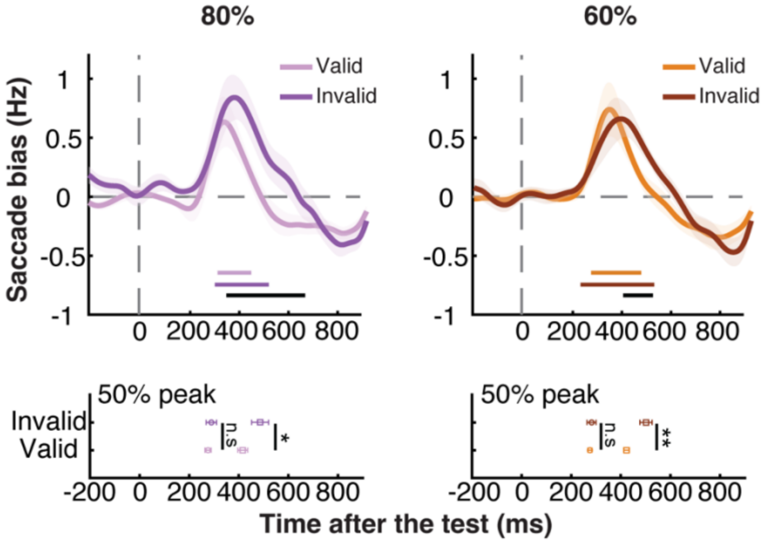


**Figure S4. Differences between the gaze bias following expected and unexpected memory tests hold robust in subsampling analysis.** Top panel: Saccade bias towards the memorized location of the tested item across conditions after the onset of the memory test in the subsampling analysis. Colored horizontal lines above the x-axes indicate significant clusters (cluster P = 0.021, < 0.001, 0.008, < 0.001, for 80%-valid, 80%-invalid, 60%-valid, 60%-invalid condition, respectively). The black horizontal lines indicate significant clusters comparing validly and invalidly cued tests (cluster P = 0.002, 0.015, for 80% and 60% condition, respectively). Bottom panel: onset and offset latency (defined by the 50% of the peak) of the spatial saccade bias following validly- and invalidly-cued memory tests in the 80% and 60% cue-reliability conditions in the subsampling analysis. Circles and squares represent onset and offset latency, the light and dark colors represent validly- and invalidly-cued memory tests. Shading and error bars indicate ±1SEM. *, **, n.s represent significance level $p < 0.05$ (0.041 for 80% cue-reliability condition), $p < 0.01$ (0.003 for 60% cue-reliability condition), and not significant, respectively.
